# Microbiota-derived pantothenic acid rapidly restores colonic mucus barrier after antibiotic-induced dysbiosis

**DOI:** 10.64898/2026.09.25.754384

**Authors:** Kota Uchiyama, Riho Furukawa, Eiji Miyauchi, Yuka Maejima, Doshun Ito, Kento Omotegawa, Riko Sawase, Chihiro Mogi, Runa Aoyagi, Tsukasa Oda, Takayuki Saitoh, Nobuo Sasaki

**Affiliations:** The Laboratory for Mucosal Ecosystem Design, Institute for Molecular and Cellular Regulation, Gunma University, Maebashi, Gunma 371-8512, Japan; Department of Laboratory Sciences, Graduate School of Health Sciences, Gunma University, 3-39-22 Showa-machi, Maebashi, Gunma 371-8511, Japan

**Author notes:** These authors contributed equally to this work (Kota Uchiyama, Riho Furukawa, Eiji Miyauchi, Yuka Maejima).

## Abstract

The intestinal mucus layer forms the primary physical barrier separating the host epithelium from the gut microbiota. Antibiotic-induced dysbiosis compromises this barrier, yet the microbial metabolites and host mechanisms responsible for its rapid restoration remain poorly understood. Here, we identify pantothenic acid (vitamin B5) as a microbial metabolite that rapidly restores colonic mucus integrity following antibiotic administration. Integrated 16S rRNA gene sequencing and metabolomic profiling of murine feces revealed a positive association between *Muribaculaceae* abundance, fecal pantothenic acid level, and colonic mucus thickness. Among four isolated *Muribaculaceae* species, only *Sangeribacter muris* produced pantothenic acid. An exogenous pantothenic acid supplement quickly restored the thin mucus layer. This rapid response occurred without detectable transcriptional activation. This response did not recur in isolated colonic epithelium or colonic organoids, indicating that pantothenic acid acts through a non-epithelial, non-autonomous mechanism. Together, these findings identify a previously unrecognized microbe-metabolite-host signaling pathway that rapidly regulates colonic mucus barrier homeostasis and reveal a local barrier-protective function of microbiota-derived pantothenic acid.

## Introduction

The mucus layer is a dynamic, continuously replenished barrier that separates the host from the dense microbial community of the gut(1, 2). This barrier is composed mainly of highly glycosylated mucins secreted by goblet cells and is essential for maintaining intestinal homeostasis by limiting microbial access to the epithelial surface(3-6). Disruption of the mucus layer compromises barrier integrity and increases susceptibility to enteric infection and inflammatory disease(7-9). Broad-spectrum antibiotics rapidly perturb the gut microbiota and are accompanied by pronounced thinning of the colonic mucus layer(10-12), yet the microbial factors required to maintain or restore this barrier remain poorly understood.

Gut microorganisms shape intestinal physiology by producing of diverse bioactive metabolites. Short-chain fatty acids (SCFAs) are among the best-characterized microbial metabolites that regulate epithelial function, immune homeostasis, and barrier integrity(13). More recently, microbiota-derived vitamins have emerged as additional mediators of host physiology. Pantothenic acid (vitamin B5), an essential precursor of coenzyme A, has recently been shown to alleviate metabolic syndrome through microbiota-dependent mechanisms(14). Whether microbiota-derived pantothenic acid also contributes to local regulation of the intestinal mucus barrier, particularly during its rapid restoration following dysbiosis, remains unknown.

Here, we investigated how microbiota-derived metabolites contribute to the restoration of the colonic mucus barrier following antibiotic-induced dysbiosis. By integrating microbiome profiling, metabolomics, and bacterial isolation, we identified pantothenic acid as a microbial metabolite associated with mucus layer integrity and *S. muris* as a producer of this vitamin. We further demonstrate that pantothenic acid rapidly restores the depleted mucus layer in vivo through a mechanism independent of *de novo* transcription and not reproduced in isolated epithelial cells or intestinal organoids, suggesting the involvement of non-epithelial intermediary cells. Taken together, these findings identify a previously unrecognized microbe-metabolite-host axis that rapidly regulates colonic mucus barrier homeostasis.

## Results

### Vancomycin and ampicillin selectively induce colonic mucus layer depletion

To determine the effects of antibiotic treatment on the colonic mucus barrier, we administered a cocktail of vancomycin, neomycin, ampicillin, and metronidazole in their drinking water for up to four weeks. To longitudinally track mucus dynamics in the same animals(15), fecal samples were collected at 1, 2, 3, and 4 weeks post-treatment, and the thickness of the mucin layer encapsulating the feces was quantified using Alcian blue staining (**Fig. 1A and B**). Control mice maintained a stable fecal mucus layer with an average thickness of approximately 20 μm throughout the experimental period (**Fig. 1B**). In contrast, mice treated with an antibiotic cocktail exhibited progressive thinning of the mucus layer that became evident after 2 weeks of administration and persisted through weeks 3 and 4 (**Fig. 1B**).

**Figure 1.**
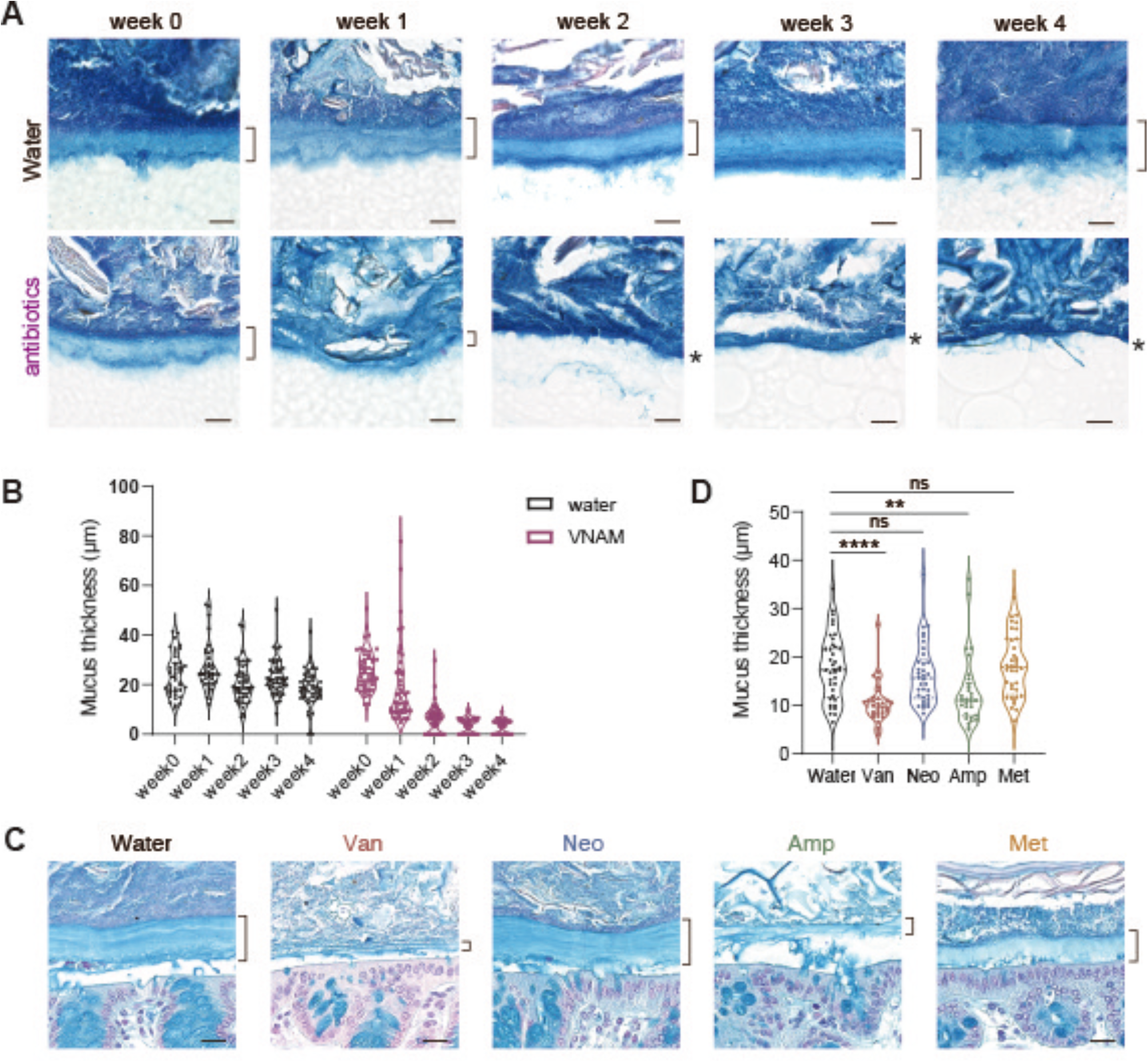
Vancomycin and Ampicillin selectively deplete the colonic mucus layer. **(A)** Representative images of Alcian blue-stained fecal pellets collected from control mice and mice treated with an antibiotic cocktail (vancomycin, neomycin, ampicillin, and metronidazole) over 4 weeks. Bracket: mucus layer thickness. Asterisk: disappearance of the mucus layer. Scale bars: 20 μm. **(B)** Time-course quantification of the mucin layer thickness encapsulating the fecal pellets from (A) at 1, 2, 3, and 4 weeks past administration. For each fecal cross-section, mucin thickness was measured at least 6 (6-12) independent locations along the circumference. Data are presented as violin plots showing the distribution of all individual measurements. Biological replicates: n = 5 mice/group for weeks 1 and 2, at weeks 3 and 4 for the control group, and n =3 for the antibiotic cocktail group at weeks 3 and 4 due to treatment-related mortality. **(C)** Representative histological images of Alcian blue-stained colonic tissue sections from mice treated individually with control, vancomycin, ampicillin, neomycin, or metronidazole for 2 weeks. Scale bars: 20 μm. **(D)** Quantification of the epithelial mucus layer thickness in the colonic tissues from (C). Data are presented as violin plots (n = 4-5 mice/group per experiment, with 2 independent experiments performed), for both (B) and (D), statistical significance compared to the control group was determined by Kruskal-Wallis test followed by Dunn’s multiple comparisons test: ^****^p<0.0001, ns= not significant.

We next asked whether a specific antibiotic could deplete the mucus layer. Mice were treated individually with ampicillin, neomycin, metronidazole, or vancomycin for 2 weeks. After treatment, intestinal tissues were collected and stained with Alcian blue to measure epithelial mucus layer thickness directly. Both ampicillin and vancomycin significantly reduced colonic mucus thickness compared to the control group, whereas neomycin and metronidazole had little or no effect (**Fig. 1C and D**). These findings indicate that a specific subset of antibiotics depletes the colonic mucus layer rather than reflecting a general consequence of antimicrobial treatment.

### Depletion of *Muribaculaceae* correlates with antibiotic-induced thinning of the colonic mucus layer

To identify specific bacterial taxa potentially involved in regulating the colonic mucin layer, we performed 16S rRNA gene amplicon sequencing of fecal microbiota from mice administered individual antibiotics for 2 weeks. Analysis of the bacterial community composition at the genus level revealed profound and antibiotic-specific shifts in the microbiota (**Fig. 2A**). Principal coordinate analysis (PCoA) based on weighted UniFrac distances demonstrated a marked effect of antibiotic treatment on overall microbial community structure (PERMANOVA, R^2^ = 0.848, *p* < 0.0001) (**Fig. 2B**). Notably, the microbiota of vancomycin- and ampicillin-treated mice, which exhibited severe depletion of the colonic mucus layer, showed pronounced separation from control mice. In contrast, the microbiota of neomycin- and metronidazole-treated mice remained closer to that of control mice, consistent with their relatively mild effects on the mucus layer.

**Figure 2.**
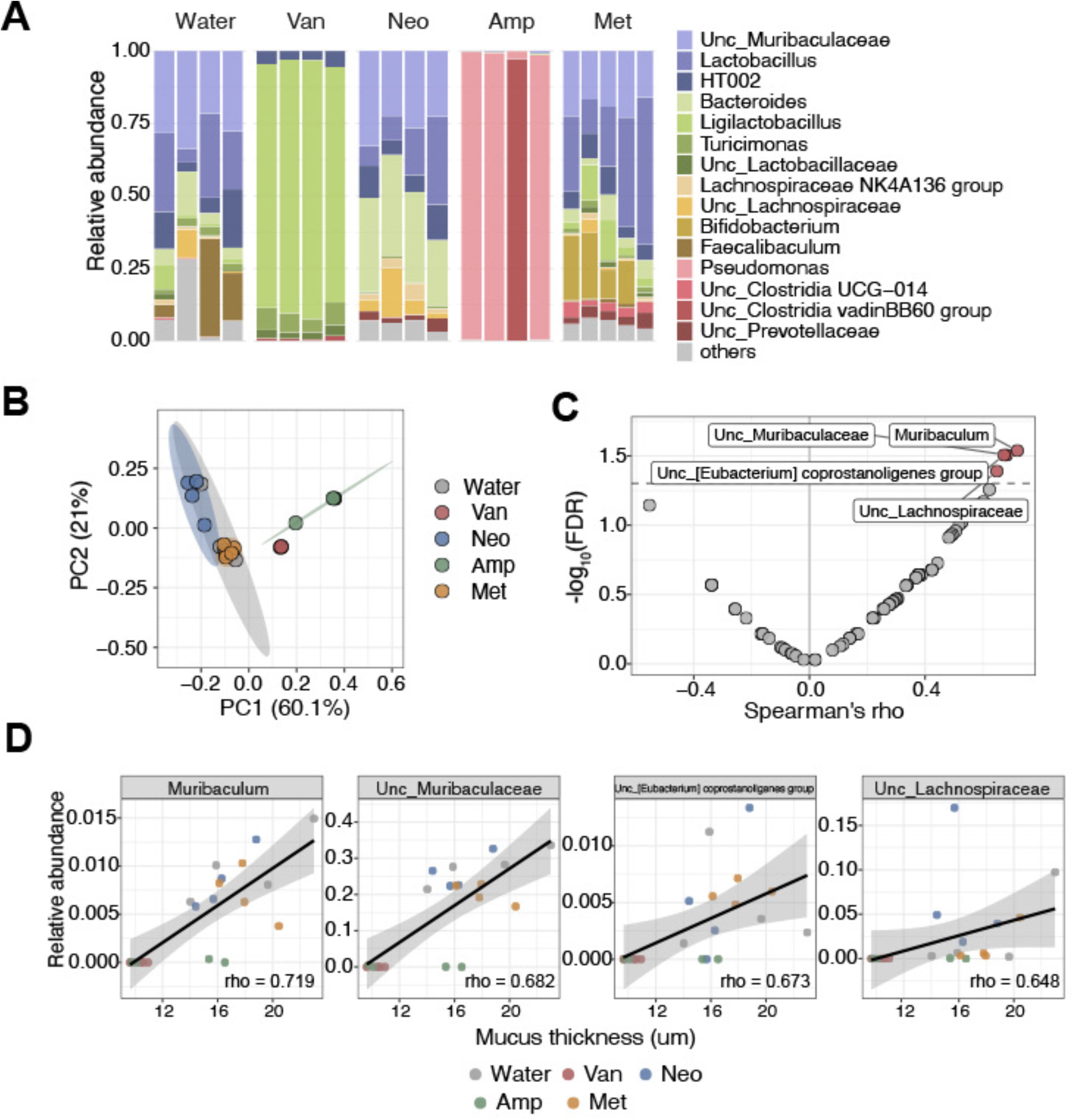
Antibiotic-induced microbiota alterations are associated with depletion of *Muribaculaceae* and thinning of the colonic mucus layer. **(A)** Genus-level composition of the fecal microbiota from individual mice treated with control, vancomycin, neomycin, ampicillin, or metronidazole for 2 weeks, as determined by 16S rRNA gene amplicon sequencing. Each bar represents one mouse. The 15 most abundant genera across all samples are shown individually, and the remaining taxa are grouped as “Others” (n = 4–5 mice/group). **(B)** Principal coordinate analysis (PCoA) of weighted UniFrac distances among the samples. Each symbol represents one mouse, and shaded areas indicate 95% confidence ellipses. The percentage of variation explained by each principal coordinate is shown on the corresponding axis. The overall effect of antibiotic treatment on microbial community structure was assessed by permutational multivariate analysis of variance (PERMANOVA) with 9,999 permutations (R^2^ = 0.848, p < 0.0001). **(C)** Volcano plot showing the correlations between the relative abundance of individual bacterial genera and colonic mucus thickness measured in Fig. 1D. Correlations were assessed by Spearman’s rank correlation with Benjamini-Hochberg correction for multiple comparisons. Each point represents one genus. The dashed line indicates a false discovery rate of 0.05, and genera meeting this threshold are labeled (n = 20 mice; one mouse in the metronidazole group was excluded because mucus thickness was not measured). **(D)** Correlations between the relative abundance of *Muribaculum* (left), Unclassified *Muribaculaceae* (second from left), unclassified *[Eubacterium] coprostanoligenes* group (third from left), and unclassified *Lachnospiraceae* (right) and colonic mucus thickness. Each symbol represents one mouse and is colored by treatment group. Lines indicate linear regression fits with 95% confidence intervals (n = 20 mice, as in (C)). Spearman’s rho is shown in each panel.

We next examined whether the relative abundance of individual bacterial taxa was associated with variation in mucus layer thickness measured in the same mice by performing Spearman’s rank correlation analysis. Four taxa, *Muribaculum*, unclassified *Muribaculaceae*, the unclassified *[Eubacterium] coprostanoligenes* group, and unclassified *Lachnospiraceae*, showed significant positive correlations with mucus layer thickness after correction for multiple testing (adjusted *p* < 0.05) (**Fig. 2C**). Notably, unclassified *Muribaculaceae* was by far the most abundance of these taxa, accounting for approximately 20% of the fecal microbiota, compared with approximately 0.5% for Muribaculum, or the unclassified *[Eubacterium] coprostanoligenes* group, and 2% for unclassified *Lachnospiraceae* (**Fig. 2D**). Its relative abundance remained largely stable following neomycin or metronidazole treatment but was markedly depleted following vancomycin or ampicillin treatment (**Fig. 2A**).

Collectively, these findings identified several bacterial taxa associated with maintenance of the colonic mucus layer. Given its substantially higher relative abundance and its marked depletion following the antibiotics that most strongly reduce colonic mucin levels, we focused subsequent investigation on unclassified *Muribaculaceae*.

### Metabolomic profiling identifies pantothenic acid depleted by mucus-disrupting antibiotics

To investigate metabolic changes associated with antibiotic-induced disruption of the colonic mucus layer, we performed metabolomic profiling of fecal samples from mice treated with the individual antibiotics. Hierarchical clustering of metabolite abundance profiles revealed two distinct clusters that closely reflected the phenotypic effects of antibiotic treatment (**Fig. 3A**). Cluster 1 (green) comprised metabolites markedly depleted following vancomycin and ampicillin treatment, the two antibiotics that caused pronounced thinning of the colonic mucus layer. In contrast, these metabolites were relatively preserved following neomycin or metronidazole treatment, which had minimal effects on mucus thickness

Notably, Cluster 1 included propionic acid and butyric acid, SCFAs previously implicated in regulating the intestinal mucus barrier(13), providing an independent metabolic correlate of the mucus phenotype (**Fig. 3A, Blue arrows)**. Interestingly, pantothenic acid was also markedly depleted following vancomycin and ampicillin treatment and clustered with propionic acid and other metabolites reduced by these antibiotics (**Fig. 3A, Red arrow**). Recent work has implicated pantothenic acid in regulating intestinal mucin production, raising the possibility that its depletion may contribute to the mucus phenotype observed following antibiotic treatment. Because our microbiome analysis identified unclassified *Muribaculaceae* as a bacterial taxon strongly associated with colonic mucus thickness (**Fig. 2D**), we asked whether unclassified *Muribaculaceae* could serve as a microbial source of luminal pantothenic acid.

**Figure 3.**
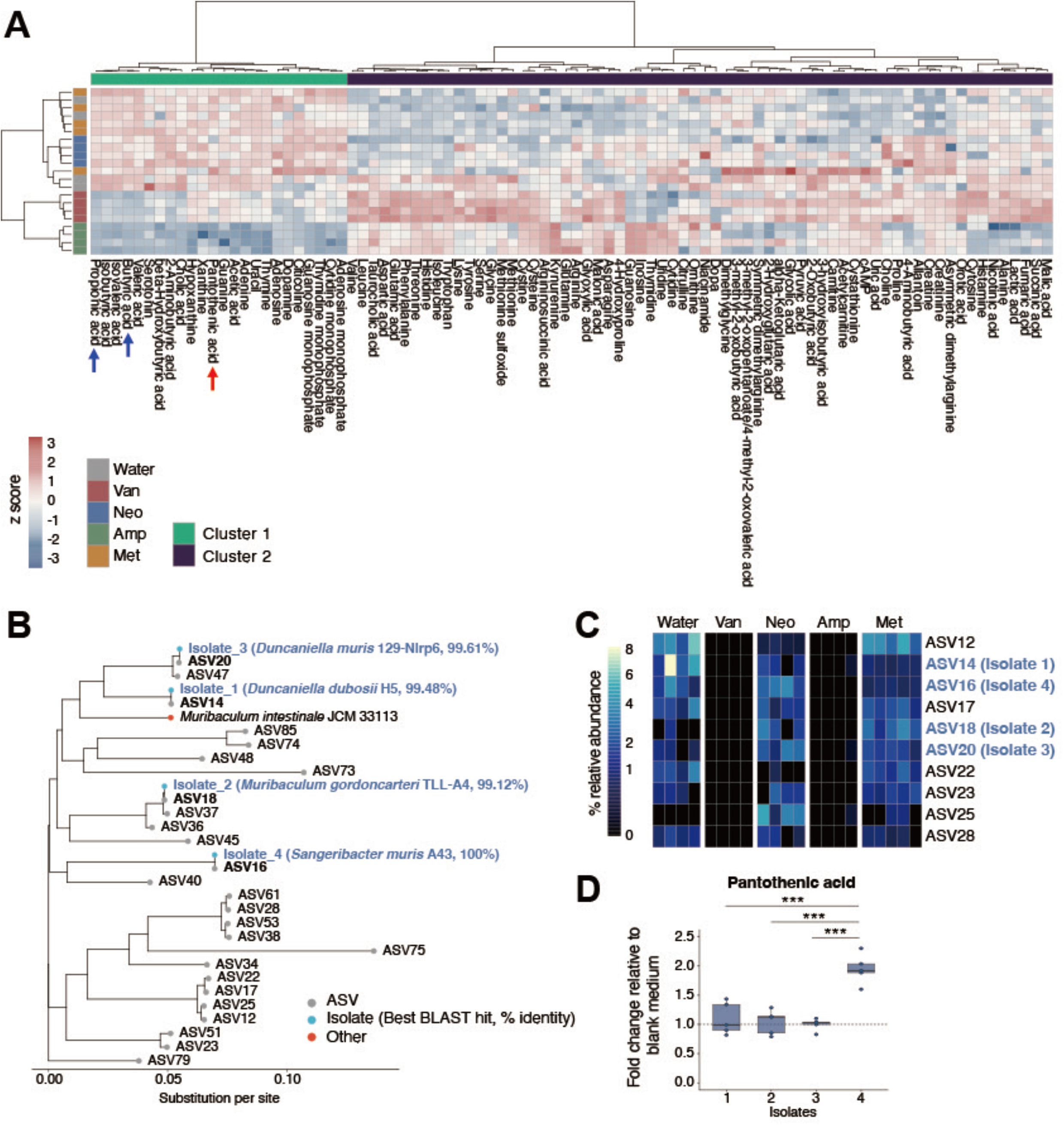
Metabolomics profiling identifies *Sangeribacter muris* as a specific microbial source of luminal pantothenic acid depleted by mucus-disrupting antibiotics. **(A)** Heatmap with hierarchical clustering of metabolomic profiles from the feces of mice treated with water only (control), vancomycin, neomycin, ampicillin, or metronidazole. Metabolites are grouped into two clusters: Cluster 1 (green) and Cluster 2 (purple). Cluster 1 represents metabolites markedly decreased by vancomycin and ampicillin. The blue and red arrows highlight SCFAs and pantothenic acid, respectively (n = 4 mice/group, except n = 5 for the metronidazole group) **(B)** Phylogenetic tree showing the relationship between the predominant unclassified *Muribaculaceae* ASVs detected by 16S rRNA gene sequencing and four newly isolated strains. For each isolate, the closest match in the NCBI 16S ribosomal RNA database and its percentage identity are shown in parentheses, and the ASV with the corresponding sequence is indicated in bold. **(C)** Heatmap showing the relative abundance of the 10 most abundant of the 26 ASVs shown in (B) in the feces of individual mice. ASVs corresponding to isolates 1–4 (ASV14, ASV18, ASV20, and ASV16) are indicated. **(D)** Quantification of pantothenic acid production in vitro. Pantothenic acid concentrations in culture supernatants from the four *Muribaculaceae* isolates were measured by LC–MS/MS and normalized to the blank medium control. Statistical significance among groups was determined by the one-way ANOVA test followed by Tukey’s multiple comparison test; ^***^*p* < 0.001

### Isolation of unclassified *Muribaculaceae* strains identifies a bacterial source of luminal pantothenic acid

To determine whether unclassified members of the family *Muribaculaceae* produce pantothenic acid, we isolated bacterial strains from mouse feces. We matched the isolates to the predominant unclassified *Muribaculaceae* amplicon sequence variants (ASVs) determined by 16S rRNA gene sequencing. Phylogenetic analysis of the cultured isolates and the unclassified *Muribaculaceae* ASVs showed that isolates 1, 2, 3, and 4 corresponded to ASV14, ASV18, ASV20, and ASV16, respectively (**Fig. 3B**). BLAST searches against 16S rRNA gene sequences deposited in the NCBI database identified *Duncaniella dubosii* H5 as the best-matching strain for isolate 1 (99.48% sequence identity), *Muribaculum gordoncarteri* TLL-A4 for isolate 2 (99.12%), *Duncaniella muris* 129-Nlrp6 for isolate 3 (99.61%), and *Sangeribacter muris* A43 for isolate 4 (100%).

We next examined the relative abundance of the ASVs corresponding to the cultured isolates in the fecal microbiota. ASV14, ASV20, and ASV16, corresponding to isolates 1, 3, and 4, respectively, were abundant in the control water-only drinking mice. In contrast, ASV18, corresponding to isolate 2, was comparatively less abundant (**Fig. 3C**). Notably, the three abundant ASVs (ASV14, 20, and 16) were markedly depleted following vancomycin and ampicillin treatment, paralleling the overall depletion of unclassified *Muribaculaceae* at the genus level observed in our microbiome profiles (**Fig. 2**). Thus, three of the cultured isolates represented abundant *Muribaculaceae* populations that were depleted by the mucus-disrupting antibiotics.

We then assessed the capacity of the four Muribaculaceae isolates to produce pantothenic acid *in vitro*. LC-MS/MS analysis showed that pantothenic acid concentrations were elevated in culture supernatants of isolate 4 (ASV16), whereas those in culture supernatants of isolates 1-3 were comparable to those measured in the uninoculated medium control (**Fig. 3D)**. Taken together, these findings suggest that the *Muribaculaceae* population corresponding to ASV16 may serve as a microbial source of luminal pantothenic acid and thereby contribute to the maintenance of the colonic mucus layer.

### Pantothenic acid rapidly restores the colonic mucus layer through an indirect mechanism

Having identified *S. muris* as a microbial source of luminal pantothenic acid that is depleted after antibiotic treatment, we next investigated whether replenishment of pantothenic acid could functionally restore the antibiotic-induced loss of the colonic mucus layer. We administered pantothenic acid intrarectally to antibiotic-treated mice with an attenuated colonic mucus layer (**Fig. 4A**). In vehicle-treated mice, the colonic mucus layer remained markedly thinner than that of untreated control mice. In contrast, intrarectal administration of pantothenic acid at 0.2, 2.0, or 20 μM restored the colonic mucus layer (**Fig. 4B and C**). Notably, this restoration occurred rapidly, with a significant increase in mucin levels detected as early as 45 min after pantothenic acid administration (**Fig. 4B and C**)

A previous study reported that long-term pantothenic acid intervention over 12 weeks restores intestinal barrier function through epigenetic changes that upregulate *MUC2* expression(14). The rapid response observed in our model therefore raised the possibility that pantothenic acid may also regulate the mucin layer through a mechanism distinct from transcriptional control, such as acute stimulation of mucin release from goblet cells. To test this possibility and determine whether pantothenic acid directly stimulates the intestinal epithelium, we first used *in vitro* colonic organoids, a defined epithelial model devoid of the surrounding lamina propria(16). Based on the concentration used in a previous study examining the effects of pantothenic acid on MUC2 expression in intestinal organoids(14), we treated colonic organoids with pantothenic acid at 320 μM. Pantothenic acid treatment did not significantly alter *MUC2* transcript levels (**Fig. 4D**). We next assessed whether pantothenic acid directly stimulates MUC2 protein secretion by measuring MUC2 in the culture supernatant of a monolayer organoid model by ELISA. Surprisingly, pantothenic acid treatment did not increase MUC2 release (**Fig. 4E**). To confirm this finding in a primary tissue context, we used an *ex vivo* model of isolated mouse colonic crypts, stripped the intestinal epithelial cells away from the underlying lamina propria, and treated those directly with pantothenic acid. Consistent with the organoid data, no changes in MUC2 secretion were observed in the isolated epithelium (**Fig. 4F**).

**Figure 4.**
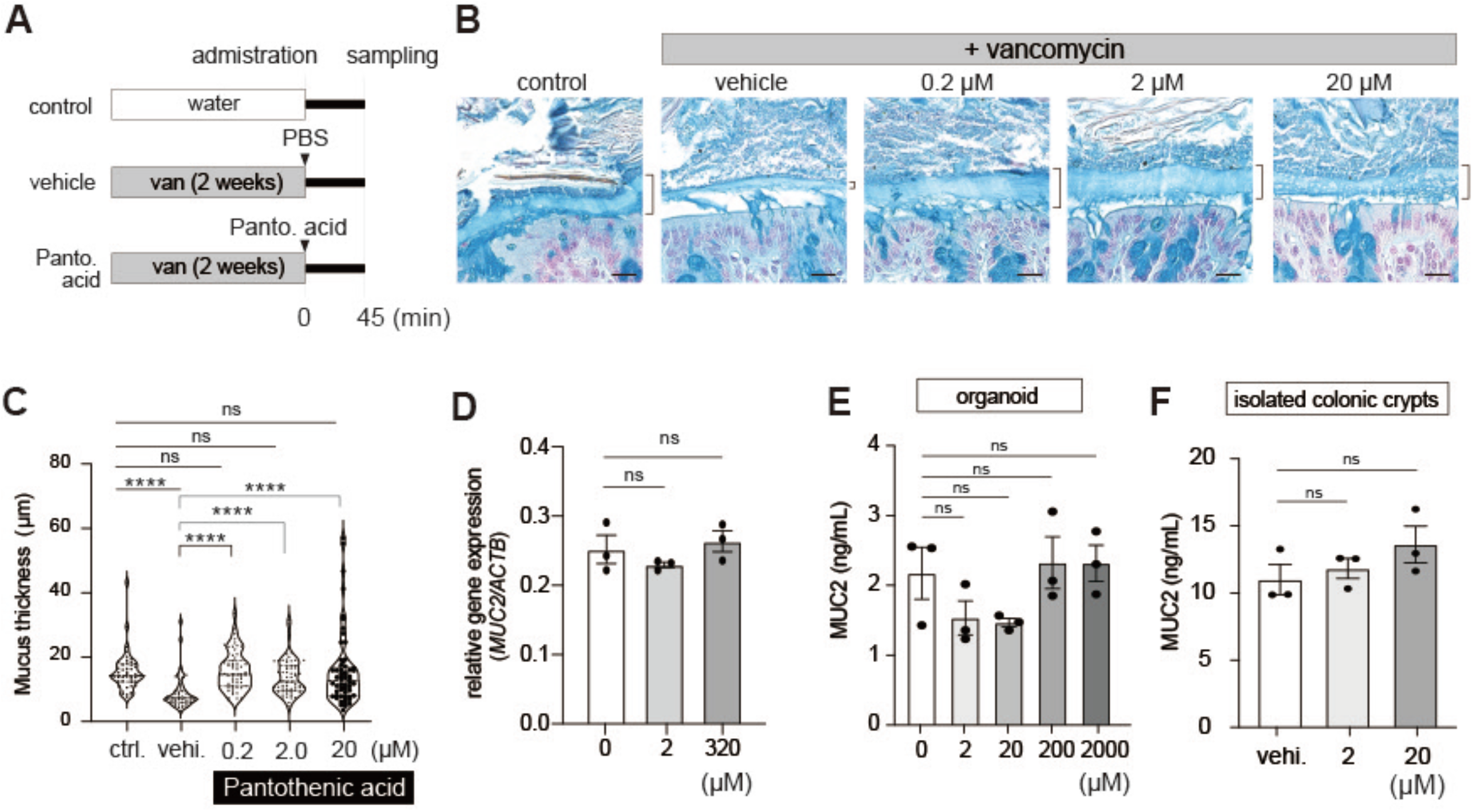
Intrarectal pantothenic acid rapidly restores the colonic mucus layer through an indirect, non-epithelial mechanism. **(A)** Experimental schema of the *in vivo* rescue trial. Mice were treated with vancomycin in drinking water for 2 weeks to deplete mucus layer, followed by intrarectal administration of vehicle or pantothenic acid (0.2, 2.0, or 20 μM). Colonic tissues were harvested 45 min post-administration. **(B)** Representative histological images of Alcian blue-stained colonic tissue sections from control mice, vancomycin-treated mice receiving vehicle, and intrarectal pantothenic acid at indicated concentrations, 0.2, 2.0, or 20 μM, respectively. Scale bars: 20 μm. **(C)** Quantification of colonic epithelial mucus layer thickness from (B). Data are presented as violin plots showing the data distribution (n = 10 mice/group, pooled from two independent experiments with n = 5 mice/group per experiment). Statistical significance compared to the control or vancomycin+vehicle group was determined by the Kruskal-Wallis test followed by Dunn’s multiple comparison test; ns = not significant, ^****^*p* < 0.0001. **(D)** Relative mRNA expression of *MUC2* in colonic organoids treated with pantothenic acid at the indicated concentration, determined by RT-qPCR and normalized to β-actin (n = 3 independent experiments). Data are presented as mean ± SEM. **(E, F)** ELISA quantification of MUC2 protein secretion in culture supernatants from a monolayer of colonic epithelium derived from organoids (E) or from isolated primary murine colonic crypts (F) treated with vehicle or indicated pantothenic acid concentrations. n = 3 independent wells for (E), n = 3 mice/group for (F). Statistical significance compared to the 0 μM (D, E) group or the vehicle control (F) was determined by the Kruskal-Wallis test followed by Dunn’s multiple comparison test.

## Discussion

The intestinal mucus layer is a highly dynamic barrier that requires constant replenishment and rapidly adapts to changes in the intestinal environment. Although the gut microbiota has been recognized as an important regulator of mucus homeostasis, the microbial factors that support rapid restoration of the mucus layer following microbiota disruption have remained elusive. In this study, we identified microbiota-derived pantothenic acid as a metabolite associated with colonic mucus integrity. We also showed that restoring pantothenic acid rapidly recovers the mucus layer following antibiotic-induced dysbiosis. Our findings further reveal that this acute response differs mechanistically from the previously described effects of prolonged pantothenic acid exposure on intestinal barrier function and *MUC2* expression(14).

Recent work by Luo *et al*. reported that pantothenic acid can directly influence intestinal epithelial cells to induce epigenetic changes, thereby upregulating transcription of the junctional molecules *ZO-1, Claudin-1*, and *MUC2*(14). This mechanism is consistent with a relatively slow remodeling of the epithelial mucin barrier program. In contrast, we observed a significant restoration of the colonic mucus layer within 45 min of local pantothenic acid administration. Such a rapid response is unlikely to depend on substantial changes in gene transcription and instead suggests that pantothenic acid can regulate mucus homeostasis through an additional, more immediate mechanism. Consistent with this interpretation, pantothenic acid did not increase *MUC2* transcription levels in colonic monolayered organoids. Furthermore, both models of *in vitro* purified epithelium organoids and *ex vivo* isolated colonic crypts showed no direct epithelial response to pantothenic acid at the secretory level. Thus, pantothenic acid appears to regulate the mucus barrier through temporally distinct mechanisms, with a rapid response that is mechanistically separable from the previously described transcriptional regulation via an epigenetic pathway.

Our data indicate that the rapid mucin release relies entirely on indirect signaling through non-epithelial intermediary cells in the underlying lamina propria. Mucin secretion from goblet cells is not driven solely by intrinsic second-messenger cascades, such as direct calcium influx(6, 17-19), but is heavily modulated by shared stimulation and cross-talk with immune and stromal populations(20-22). For example, TNF-α and IL-1β stimulate mucin secretion and increase the expression of gel-forming mucins, including MUC2 (23, 24). Other cytokines, including type 2 cytokines such as IL-4 and IL-13, promote goblet cell differentiation and mucin gene expression through STAT6-dependent pathways, whereas IL-22 regulates mucin production and goblet cell responses through STAT3-dependent signaling(25-28). Stromal cells can also contribute to mucus regulation through cytokine-mediated epithelial-stromal crosstalk, while innate lymphoid cells provide additional signals that influence goblet cell function(29, 30). Thus, a complex network of epithelial-intrinsic and tissue-derived signals regulates the intestinal mucus, not epithelial cell-autonomous mechanisms alone. Our finding that pantothenic acid rapidly restores the mucus layer *in vivo* but fails to stimulate *MUC2* expression or mucin secretion in isolated epithelial models suggests that pantothenic acid acts on one or more non-epithelial populations to generate secondary signals that promote acute mucus release or accumulation. Identification of the pantothenic acid-responsive cell population and the downstream mediators responsible for this rapid response will be an important direction for future study.

In conclusion, our study identifies a microbiota-derived pantothenic acid signal that quickly restores the colonic mucus barrier following antibiotic-induced dysbiosis. This rapid response occurs independently of direct epithelial stimulation and differs from the previously described transcriptional regulation of *MUC2*. These findings reveal an additional layer of gut bacteria-host communication in which a microbial metabolite can rapidly influence mucus barrier homeostasis through the intestinal mucosal microenvironment.

## Materials and Methods

### Animals and antibiotic treatment

Male Jcl/ICR mice were purchased from CLEA Japan (Tokyo, Japan) and maintained under SPF conditions. All experiments were approved by the Institutional Review Board for Animal Experiments at Gunma University (study ID: animal 24-066) and were performed according to the institutional guidelines and home office regulations. Antibiotics (ampicillin (1.0 g/L; Sigma-Aldrich), vancomycin (0.5 g/L; FUJIFILM Wako), metronidazole (1.0 g/L; FUJIFILM Wako), neomycin sulfate (1.0 g/L; Sigma-Aldrich) were dissolved in drinking water to the final concentration and filtered through a 0.22 μm filter (MERCK). These four types of combination antibiotics, or each antibiotic individually, were administered to mice via the free-drinking method. Bottles of drinking water containing antibiotics were changed twice a week to maintain their effectiveness.

### Histology and measurement of the mucus thickness

To preserve the intestinal mucus layer, collected fecal pellets (from the 4 -week time course) and complete colonic tissues (from the 2-week individual antibiotic experiment) were immediately fixed in Methanol-Carnoy’s fixative. Samples were then dehydrated, paraffin-embedded, and cut into 5 μm-thick cross-sections. Following deparaffinization and rehydration, samples were stained with Alcian blue (Fujifilm Wako) and Nuclear Fast Red (Vector Laboratories). Stained sections were visualized using a Keyence BZ-800 Fluorescence Microscope, and the thickness of the mucus layer (the blue-stained area engulfing the fecal pellet) was measured using ImageJ software. For the longitudinal fecal sample analysis, one complete cross-section of a fecal pellet was analyzed per mouse per time point. The thickness of the mucin layer adhering to the circumference of the feces was measured at 6 to12 independent, evenly spaced locations around the pellet to account for spatial variability. For colonic tissue analysis, the thickness of the inner epithelial mucus layer was measured at 6 to 12 distinct locations per mouse along the distal colon. All measurements were performed by an investigator blinded to the experimental groups.

### Bacterial DNA extraction from fecal samples

Fecal samples were collected from mice and stored at -80°C until processing. Fecal samples were weighed and thoroughly suspended in Tris-EDTA (TE) buffer, subsequently filtered through a 40-`µm cell strainer (Falcon).` Bacterial DNA was isolated by the enzymatic lysis method using lysozyme (300 mg/mL, Fujifilm Wako), purified achromopeptidase (20000 U/mL, Fujifilm Wako), and proteinase K (25 mg/mL, Merck). Genomic DNA was isolated with Phenol/chloroform/isoamyl alcohol (Nacalai Tesque) and precipitated with ethanol (Fujifilm Wako) and sodium acetate (Fujifilm Wako). DNA was then centrifuged for 15 min at 14,000 rpm at 4°C, rinsed with 70% ethanol, and resuspended in TE buffer.

### 16S rRNA gene sequencing and analysis

The V3-V4 regions of the 16S rRNA gene were amplified from fecal DNA by PCR using the 341F (5′-TCGTCGGCAGCGTCAGATGTGTATAAGAGACAGCCTACGGGNGGCWGCAG-3′) and 805R (5′-GTCTCGTGGGCTCGGAGATGTGTATAAGAGACAGGACTACHVGGGTATCTAATCC-3′) primers. The resultant amplicons were indexed using the Nextera XT Index Kit v2 (Illumina). The concentration of the pooled library was determined by quantitative PCR with the KAPA Library Quantification Kit (Roche), and the fragment size was verified using a MultiNA microchip electrophoresis system (Shimadzu). Paired-end sequencing (2 x 300 bp) was then carried out on an Illumina MiSeq platform. The resultant sequencing reads were processed using DADA2 (v1.36.0) in R (v4.5.1) according to the DADA2 tutorial pipeline(31). After quality filtering, denoising, merging of paired-end reads, and removal of chimeric sequences, amplicon sequence variants (ASVs) were inferred. Taxonomic assignment was performed using the SILVA database (v138.2)(32). ASVs assigned to mitochondria or chloroplasts were excluded from subsequent analyses. Principal coordinate analysis based on weighted UniFrac distances was performed using the phyloseq package (v1.52.0) in R. Spearman’s rank correlation coefficients between genus-level relative abundances and mucus thickness were calculated with the microbiome package (v1.30.0), and P values were adjusted for multiple comparisons using the Benjamini-Hochberg method. Data visualization was performed using the ggplot2 (v4.0.3) and pheatmap (v1.0.13) packages.

### Preparation of inulin-supplemented blood agar

The basal agar contained 13.0 g soy peptone (Nacalai Tesque), 18.0 g proteose peptone (Gibco), 5.0 g yeast extract (Gibco), 2.2 g meat extract (Nacalai Tesque), 2.5 g potassium dihydrogen phosphate (KH_2_PO_4_) (Nacalai Tesque), 3.0 g sodium chloride(Fujifilm Wako), 0.3 g L-cysteine hydrochloride(Wako), and 15.0 g agar (Fujifilm Wako) in 800 mL Milli-Q water. Hemin (Alfa Aesar) solution was prepared by dissolving 10.0 mg hemin in 1.00 mL 1 N NaOH and bringing the volume to 25.0 mL with Milli-Q water. Vitamin K_3_ (Nacalai Tesque) solution was prepared by dissolving 5 mg vitamin K_3_ in 1 mL ethanol and bringing the volume to 25.0 mL with Milli-Q water. Add both solutions to the basal medium before autoclaving. After autoclaving, dissolve 8.0 g inulin (Tokyo Chemical Industry) in 100 mL Milli-Q water at 50–70ºC, then aseptically add 50 mL defibrinated sheep blood to yield approximately 1 L of medium. [Add manufacturers/catalog numbers for the remaining key reagents and state the autoclave conditions and temperature at which blood was added.]

### Isolation and cultivation of candidate bacteria

A 10-week-old Jcl:ICR mouse obtained from CLEA Japan, Inc. (Ishibe Breeding Facility, Shiga, Japan) was euthanized by cervical dislocation immediately after arrival. The colon was excised and immediately transferred to the anaerobic chamber containing 80% N_2_, 10% CO_2_, and 10% H_2_. Inside the chamber, collect a fecal pellet from the colon into a 1.5-mL tube. The sample was suspended in 0.5 mL PBS, vortexed thoroughly, and allowed to stand for approximately 1 min to sediment large debris. A 50-µL aliquot of the supernatant was transferred to 450 µL PBS, and 10-fold serial dilutions were prepared to 10^−5^. Aliquots (100 µL) of each dilution were spread onto inulin-supplemented blood agar and incubated anaerobically at 37ºC for 4-5 days. *Muribaculum intestinale* JCM33113 was cultured in parallel as controls where applicable. Distinct colonies were picked and inoculated into 200 µL starch-containing broth in 96-well plates, followed by incubation anaerobically at 37ºC for 3 days. After growth, 100 µL of each culture was transferred to a separate PCR plate for crude DNA preparation. The remaining 100 µL was mixed with 100 µL of 40% glycerol (final concentration, 20%), the plate perimeter was sealed with vinyl tape, and stocks were stored at -80ºC.

### Crude DNA preparation and qPCR screening

Crude DNA templates were prepared from the 100-µL culture aliquots by three freeze–thaw cycles between -80ºC and 95ºC. The resulting bacterial lysates were diluted 1:100 in TE buffer before PCR analysis. Candidate isolates were screened by qPCR using a Lactobacillus-specific primer set for negative selection and an uncultured *Muribaculaceae*-targeting primer set for positive selection. Each 10-µL reaction contained 5.0 µL PowerUp SYBR Green Master Mix, 0.5 µL of a 10 µM primer mixture, 3.5 µL nuclease-free water, and 1.0 µL of diluted lysate. Amplification was performed at 50ºC for 2 min and 95ºC for 2 min, followed by 40 cycles of 95 for 15 s, 62ºC for 30 s, and 72ºC for 1 min.

### 16S rRNA gene amplification and Sanger sequencing

The nearly full-length 16S rRNA gene was amplified from the 1:100-diluted lysate using primers 27F (5’-AGAGTTTGATCMTGGCTCAG-3’) and 1492R (5’-GGYTACCTTGTTACGACTT-3’).

Each 50-µL reaction contained 4.0 µL dNTP mixture, 5.0 µL 10× reaction buffer, 0.25 µL Ex Taq DNA polymerase, 3.0 µL of a 5 µM 27F/1492R primer mixture, 36.75 µL nuclease-free water, and 1.0 µL diluted lysate. PCR was performed with an initial denaturation at 94 ºC for 1 min; 30 cycles of 94 ºC for 30 s, 53 ºC for 30 s, and 72 ºC for 2 min; and a final extension at 72 ºC for 7 min. Amplicons were verified by electrophoresis on a 2% agarose gel and submitted to Eurofins Genomics for capillary Sanger sequencing using 27F, 334F (ACTCCTACGGGAGGCAGCAGT), 515F (GTGCCAGCMGCCGCGGTAA), and 1492R. For each Uni334F sequencing submission described in the record, 1 µL purified PCR product and 1 µL 10 µM 334F primer were combined with 19 µL nuclease-free water and held at 4 ºC until collection.

### Sequence processing and taxonomic identification

Raw Sanger chromatograms were inspected, low-quality terminal regions and ambiguous base calls were removed, and overlapping reads were assembled into a consensus sequence using ApE (v3.1.9). The consensus sequence—or the quality-trimmed partial read when only one sequencing primer was used—was compared with reference sequences using BLASTn against the NCBI 16S ribosomal RNA sequences (Bacteria and Archaea) database. Taxonomic identification was based on the best-supported matches after considering percentage identity, query coverage, E-value, and consistency among the top hits. Results were reported as the closest 16S rRNA gene match unless species-level assignment was supported by the available sequence resolution.

### Sample preparation for metabolome analysis

To analyze fecal metabolites, a fecal sample was suspended in 500 µL of methanol with the internal standards (8 µM 2-Ethylbutyric acid (Sigma-Aldrich), 10 µM 2-Morpholinoethanesulfonic acid (DOJINDO), 10 µM L-Methionine sulfone (Thermo Fisher Scientific). The mixture was combined with four 3-mm zirconia beads and about 0.1 g of 0.1-mm zirconia/silica beads (TOMY) and shaken vigorously for 5 min using a TissueLyser II (QIAGEN).

For bacterial culture supernatant analysis, each bacterial strain was first grown overnight. The overnight culture was mixed with fresh medium at a 1:1 ratio and incubated for 4 h as a pre-culture. The culture was then diluted to an OD_600_ of 0.1 in fresh medium and incubated while monitoring bacterial growth. Culture supernatants were collected when each strain reached the stationary phase. Because growth kinetics differed among strains, collection times were adjusted for each strain. Supernatants were collected at 9 h for *M. gordoncarteri*, 10 h for *D. muris*, 21 h for *S. muris*, and 28 h for *D. dubosii* after inoculation at an OD_600_ of 0.1. The suspension was centrifuged at 20,000 x g for 10 min at 4ºC. 100 µL of the supernatant was mixed with 400 µL of methanol with the internal standards and centrifuged at 20000 x g for 15 min at 4ºC. The supernatant was applied to the MonoSpin phospholipid column (GL Sciences) and centrifuged at 3000 x g for 30 seconds at 4ºC. The eluate was stored at -80ºC until use. For short-chain fatty acids (SCFAs), sample derivatization was performed as follows. 25 µL each of three reagents, 50 mM 1-ethyl-3-(3-dimethylaminopropyl) carbodiimide (EDC; Sigma-Aldrich), 50 mM 3-nitrophenylhydrazine (3-NPH; Sigma-Aldrich), and 7.5% pyridine (Fujifilm Wako Chemical), all dissolved in 75% (v/v) MeOH, were added to 50 µL of the sample. The mixture was vortexed for 30 min at room temperature in the dark to allow the derivatization reaction to proceed. After derivatization, an equal volume (125 µL) of 2% formic acid/75% methanol solution was added to quench the reaction. For primary metabolites, 300 µL of methanol were added to 200 µL of the samples and mixed with 250 µL of Milli-Q water and 500 µL of chloroform, vortexed for 3 min, and centrifuged at 20000 x *g* for 10 min at 4ºC. Subsequently, 300 µL of the aqueous layer was centrifugally filtered through a 5-kDa cutoff filter (UltrafreeMC-PLHCC, Merck) at 9100 x *g* for 3 h at 4ºC. The filtrate was centrifuged to concentrate and dissolved in 50 µL Milli-Q water immediately before LC-MS/MS analysis.

### LC-MS/MS analysis

Metabolites were assessed using liquid chromatography coupled to a triple quadrupole mass spectrometer (LC-MS/MS; LC-MS-8050 system, Shimadzu). LC was performed using a Mastro2 C18 column (2.0 mm internal diameter × 150 mm length, 3 µm particle size; Shimadzu Corporation) for SCFAs or a Discovery HS F5-3 HPLC column (2.1 mm internal diameter × 150 mm length, 3 µm particle size; Sigma-Aldrich) for primary metabolites. The relative levels of metabolites were determined using the Method Package for SCFAs or the Method Package for Primary Metabolites (Shimadzu). The peak area for each metabolite was normalized by the peak area of the internal standard. We processed the fecal metabolomic data using MetaboAnalyst (v6.0). Metabolites detected in fewer than 50% of samples were excluded, followed by removal of features with low variance using the default variance-based filtering criterion. Missing values were imputed with one-fifth of the minimum positive value for each metabolite. The resulting data were log10-transformed, mean-centered, and scaled to unit variance. Principal component analysis was performed using MetaboAnalyst, and the results were visualized using the ggplot2 package. Hierarchical clustering of metabolites and samples was performed using Euclidean distance and Ward’s method, and a heatmap was generated using the pheatmap package.

### Human Colonic Organoid

We used patient samples obtained at Gunma University Hospital, Gunma, Japan. We collected tissues from patients undergoing elective surgery. The establishment of patient-derived organoids and their downstream study were approved by the Gunma University Ethical Review Board for Medical Research Involving Human Subjects (Approval ID: HS2022-054 and HS2026-151). Written informed consent was obtained from all patients. Normal human colonic tissues were obtained from surgical specimens. Normal intestinal tissues were cut into 5-10 mm3 fragments using forceps and washed with cold PBS. The tissues were treated with 5 mM EDTA (Thermo Fisher Scientific) for 45 min at 4ºC with gentle rocking, and the intestinal epithelial crypts were released by pipetting. The isolated crypts were embedded into Matrigel (Corning) domes and overlaid with the organoid medium as follows: Advanced Dulbecco’s Modified Eagle’s Medium/F12 (Thermo Fisher Scientific) supplemented with 10 mM HEPES (Thermo Fisher Scientific), 2 mM GlutaMAX (Thermo Fisher Scientific), penicillin/streptomycin (Nacalai Tesque), B-27 supplement (Thermo Fisher Scientific), 1 mM N-acetylcystein (Sigma-Aldrich), 50 ng ml^-1^ recombinant mouse EGF (Thermo Fisher Scientific), 100 ng ml^-1^ recombinant mouse Noggin (PeproTech), 1 μg ml^-1^ recombinant human R-spondin1 (R&D), 100 ng ml^-1^ recombinant human IGF-1 (Biolegend), 50 ng ml^-1^ FGF-MAX (MBL), 500 nM A83-01 (Tocris), 50% Wnt3a conditioned medium from Wnt3a expressing 293T cells (A kindly gift from Hans Clevers).

### Monolayer Culture of Colonic Organoids and Pantothenic Treatment Assay

To establish monolayer organoid culture, human colonic organoids were maintained and processed as previously described with minor modifications (33). Briefly, 24-well Thincert culture inserts (0.4 μm pore size, Greiner Bio-One) were coated with 5% Matrigel diluted in Advanced DMEM/F12 at 37ºC for 30 min, then dried in a tissue culture hood. One day before single-cell dissociation, three-dimensional organoids cultured for 3 to 5 days were treated with 10 μM Y-27632 (FUJIFILM Wako). The organoids were then dissociated into single cells using TryPLE Express (Thermo Fisher Scientific), resuspended in organoid culture medium containing 10 μM Y-27632 at 1×10^6^ cells/mL, and seeded onto the coated Thincert inserts (200 μL per insert). Cells were maintained in Y-27632-supplemented medium for the first 2 days post-seeding. Thereafter, the culture medium was refreshed every 2 days without Y-27632 until confluent epithelial monolayers were established. For the pantothenic acid stimulation assay, confluent organoid-derived monolayers were treated with pantothenic acid at the final concentration indicated in Figure 4E and F, and incubated at 37ºC for 45 min.

### ELISA for Muc2

MUCIN 2 was measured using the MUC2 ELISA kit (Cloud-Clone Corp and Abcam) according to the manufacturer’s protocol.

### Real-Time Quantitative PCR

Total RNA was extracted from monolayered organoids using the RNeasy Mini Kit (QIAGEN), and cDNA was synthesized using ReverTra Ace (TOYOBO) according to the manufacturers’ instructions. Real-time quantitative PCR (RT-qPCR) was performed using THUNDERBIRD Next SYBR qPCR Mix (TOYOBO) on a CFX Opus 96 Real-Time PCR System (Bio-Rad). Relative gene levels were calculated using the 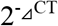 method, normalized to an internal control gene (*ACTB*).

### Quantification and Statistical Analysis

GraphPad Prism software (version 8.0 and 9.0) was used for statistical analyses. Results are shown as means ± SEM, depending on the number of samples. Groups of data were compared using one-way analysis of variance (ANOVA) with Tukey multiple comparison post hoc analysis, the Student t-test, Kruskal-Wallis test followed by Dunn’s multiple comparison test, or the Mann-Whitney U test. Differences in fecal microbial community structure among groups were evaluated by PERMANOVA based on weighted UniFrac distances (9,999 permutations; vegan v2.7.5). The difference was considered significant when the p-value was less than 0.05. All statistical details are provided in each figure legend.

## Acknowledgment

The authors thank all members of the laboratory for Mucosal Ecosystem Design for their assistance with experiments and discussion, and Dr. Yoshihiko Hagiwara (IMCR Joint Usage/Research Support Center, Gunma University) for technical assistance. The LC-MS/MS analysis was performed using an LCMS-8050 at the Core facility Management and Technical Collaboration Center (CoMTeCC) of Gunma University. The authors also thank the Mouse Facility Core of Gunma University for its assistance.

## Author contributions

K.U., R.F., Y.M., D.I., R.S., C.M., R.A., and T.O. performed the experiments and analyzed the data. K.U., E.M., and K.O. analyzed 16S rRNA gene sequencing data. K.U., R.F., and D.I. analyzed metabolome data. R.F., C.M., R.S., and R.A. performed the mouse experiment and histological analysis. Y.M. performed the organoid experiments. T.S. reviewed and edited the manuscript and participated in discussion. N.S. conceived and supervised the project and acquired funding. K.U., E.M., and N.S. wrote the manuscript. All authors discussed the results and edited the manuscript.

## Declaration of generative AI and AI-assisted technologies in the writing process

The authors used Gemini 3.1 pro from Google and Grammarly to verify grammar and improve readability during the writing and editing of this manuscript. After using these services, the authors reviewed and edited the manuscript as needed and assumed full responsibility for the content of the published article.

